# Development of a novel GWAS method to detect QTL effects interacting with the discrete and continuous population structure

**DOI:** 10.1101/2024.03.13.584913

**Authors:** Kosuke Hamazaki, Hiroyoshi Iwata, Tristan Mary-Huard

## Abstract

Although GWAS has been a key technology to identify causal genes, the current standard GWAS model still has problems that need to be solved. Among them, the population structure is one of the most severe problems when detecting QTLs in GWAS since the GWAS model is statistically confounded by effects derived from the population structure. Further, the existence of QTLs, whose effects depend on the genetic background, also affects the conventional GWAS results by causing many false positives. Although the model to detect these population-specific QTLs has already been developed, this model requires prior information on the population structure, which may only sometimes be available. Also, the previous model only assumed the situation where QTLs interact with the discrete population structure. However, target populations of GWAS often consist of genetic resources with a more continuous population structure, and there has been no model that can consider such QTLs interacting with the continuous structure. In this study, by explicitly including an interaction term between a SNP/haplotype block and the genetic background in the conventional SNP-based/haplotype block-based GWAS model, we developed two models, named SNPxGB and HBxGB, that can detect QTLs interacting with the discrete and continuous structure. Our developed models were compared to the previous models by a simulation study assuming some types of QTLs, i.e., QTLs with effects common to all the backgrounds, specific to one genetic background, and interacting with polygenes. The simulation study showed that the models assuming the same situation as the simulation settings for each QTL type were suitable for detecting the corresponding QTLs. Primarily, our second HBxGB model could detect QTLs interacting with polygenes, i.e., continuous population structure, better than the previous model utilizing the prior population structure information. Our developed models are expected to help unravel the unknown genetic architecture of many complex traits.

**Author summary:** GWAS aims at detecting candidate genes associated with a target trait via statistical testing. Since a classical GWAS starts with the constitution of a panel of individuals, usually gathered from different populations, many methods have been proposed to control the false positives in large datasets with a strong population structure. However, most methods assume the same QTL effect across populations, which is not always true in the natural biological process. One study has proposed a method to consider population-specific QTL effects by assuming marker effects depend on each subpopulation with prior information on population membership for each individual. This information on the population structure, however, may only sometimes be available, and sometimes the population structure is more continuous rather than discrete, where their methodology cannot be applied. We successfully developed two novel models that do not require prior knowledge of the population structure by explicitly including an interaction term between a SNP/haplotype block of interest and the genetic background in the conventional SNP-based/haplotype block-based GWAS model. The developed models, named SNPxGB and HBxGB, were suitable for capturing gene effects interacting with the discrete and continuous population structure, leading to the clarification of the genetic architecture of complex traits.

## Introduction

In recent years, the advent of next-generation sequencing technologies has enabled the collection of high-density whole-genome sequencing data for many species, leading to the high-throughput genotyping of large numbers of accessions at lower cost [1–3]. The genomic data obtained with these new sequencing and genotyping technologies has been utilized for various genetic analyses, including genome-wide association studies (GWAS) [4,5]. Following its first applications in human genetics [6–8], GWAS has contributed to the identification of novel genes not only in human genetics but also in plant and animal genetics and breeding [9–12], and its momentum is still increasing with the use of whole-genome sequencing technologies [13,14].

A classical GWAS starts with the constitution of a panel of individuals, usually gathered from different populations [15]. This population stratification must be accounted for in the analysis to efficiently control the Type I error rate of the quantitative trait loci (QTL) detection procedure [16,17]. Many methods have been proposed to control the false positive error rate in large datasets with strong population structure [18–20], e.g., a mixed-effects model to correct the effects of family relatedness by regarding polygenetic effects as random effects [21]. While controlling for false positive, these methods improved the QTL detection in GWAS with genetically diverse populations while controlling false positives well [14,22–24].

These conventional GWAS models and subsequent studies have implicitly assumed that the QTL effects are constant across different genetic backgrounds in a target population. However, in the natural biological process, the effects of causal genes or QTLs sometimes depend on the genetic background (S1 Fig for an example of plant height) [25–27]. These QTLs with effects depending on the genetic background are called group-specific/population-specific QTLs, whose effects may be caused by epistasis between QTLs and other loci with population-specific allele frequencies [28,29], a different pattern of linkage disequilibrium (LD) between SNPs and QTLs across populations [27], and/or population-specific mutations within QTL regions [31]. Rio et al. proposed a model to detect these population-specific QTLs [27]. Concretely, utilizing the maize inbred lines consisting of two subpopulations of Dent and Flint and the admixed population between them, they developed a model assuming that each SNP effect depends on its ancestry, i.e., Dent or Flint, and genetic background, i.e., Dent, Flint or admixed population. Applied to the analysis of flowering date in maize, the model successfully detected population-specific QTLs while controlling false positives and succeeded in disentangling population-specific QTL effects in light of the above three reasons [27].

The model proposed by Rio et al. relies on the availability of information about the genetic background of each individual and ancestral subpopulation for each marker of each admixed individual [27]. In applications where the panel under study was obtained based on a controlled multiple crossing design such as nested association mapping (NAM) [36] or multi-parent advanced generation inter-cross (MAGIC) [37] populations, or alternatively, when the panel is highly structured, the model is expected to significantly improve the QTL detection.

However, target populations of GWAS often consist of genetic resources with a more cryptic and continuous population structure rather than a discrete one, e.g., due to the presence of individuals exhibiting a wide range of admixture levels [38]. In such cases, the lack of a clear, discrete population structure could hamper the application of the conventional GWAS methods, as discussed in the context of the mixed-effects model to correct the effects of family relatedness [21,36,37]. Moreover, in analytical populations with a cryptic and continuous structure, the first reason for population-specific QTLs, i.e., epistasis between QTLs and other loci with varying allele frequencies [38], will evolve into an epistatic interaction between QTLs and polygenes since genome-wide polygenes contribute to the formation of population structure. However, there has been no model that can account for such QTLs interacting with polygenes under a continuous population structure.

In the present study, we develop two novel methods that do not require prior knowledge of the population structure (S2 Fig). Our models focus on the fact that the population-specific QTL effects can be regarded as interaction effects between QTLs and the genetic background. More specifically, in our first SNPxGB model, we explicitly include an interaction term between a SNP of interest and the discrete/continuous population structure in the conventional GWAS model of Yu et al. [21]. Further, in our second HBxGB model, we mainly aim to consider interaction effects between QTLs and the continuous population structure, which have not been considered in the previous models. To achieve this, based on a haplotype block (HB) based GWAS model that tests multiple SNPs in each HB at a time [38], we include an interaction term between an HB of interest and the continuous population structure consisting of polygenes in the HB GWAS model. By developing these two GWAS models, our ultimate goal is to detect various types of QTLs whose effects depend on the discrete and continuous population structure, which will result in unraveling the genetic architecture of agronomically important quantitative traits. The validity of the proposed models was evaluated based on a simulation study using a soybean dataset, and then, the models were also applied to real phenotypic data.

## Materials and Methods

### Materials and marker genotype data

A panel of 900 soybean accessions was sampled from the USDA Soybean Germplasm Collection (available from the SoyBase, www.soybase.org) [39]. The whole collection consists of nearly 22,000 accessions of *Glycine max* and *Glycine soja* [39,40], collected from 89 countries and genotyped using the Illumina Infinium SoySNP50K BeadChip [41]. A first data reduction was performed by filtering markers with a missing rate ≤ 0.01 and a minor allele frequency (MAF) ≥ 0.05, and by selecting accessions collected from three countries: China, Japan, and Korea, hereinafter referred to as subpopulations. In each of the three subpopulations, a set of 300 representative accessions were selected as follows: accessions within each subpopulation were clustered into 300 groups using a k-medoid procedure [42], then the 300 medoid accessions representative of each class were selected for subsequent analyses.

After selecting the 900 accessions, missing SNP data were imputed using Beagle version 5.4 [43,44]. Then, a second marker filtering for MAF (≥ 0.025) was applied only to bi-allelic sites over all accessions, leading to 38,634 biallelic SNPs. Finally, a third SNP filtering was applied within each population, resulting in 27,100, 27,786, and 28,214 SNPs with MAFs ≥ 0.05 and LD ≤ 0.95 in China, Japan, and Korea subpopulations, respectively. In the analysis, the genotypes were represented as −1 (homozygous for the reference allele), 1 (homozygous for the alternative allele), or 0 (heterozygous).

Haplotype blocks (HBs) were estimated based on marker genotype data using PLINK 1.9 [45–47]. Parameters were set to their by-default values except for “--blocks-min-maf”, which was changed to 0.025 to allow the presence of rare variants in each HB during the HB estimation. The procedure resulted in 20,089 HBs. Also, we estimated LD blocks as the blocks whose lengths were longer than HBs by using PLINK, which resulted in 12,754 LD blocks. These LD blocks were utilized to evaluate the GWAS results in this study. Here, when a SNP was not included in the original blocks estimated by PLINK, HB and LD blocks were imputed as blocks consisting of that SNP.

As for the software used in the pre-processing, the first and second data reductions for the marker filtering with missing rate and MAF were conducted by using VCFtools version 0.1.16 [48]. The third per-subpopulation filtering with MAF and LD was conducted by the MAF.cut function in the R package RAINBOWR version 0.1.31 [38] and the LD.thin function in the R package gaston version 1.5.9 [49,50]. The selection of the representative accessions was performed based on the marker genotype data consisting of 37,233 SNPs after the first marker filtering, using the “pam” function of the R package “cluster” version 2.0.9 [50]. As described above, we also used the Beagle version 5.4 [43,44] for the imputation of missing SNPs before the second marker filtering and the PLINK 1.9 [45–47] for the estimation of the HBs after the second marker filtering. Source codes for the pre-processing, including the parameters for the PLINK, are available on GitHub, as explained in the **Code availability** subsection.

### Simulation of phenotype data

First, we conducted a simulation study to evaluate the validity of the proposed models described as below. To simulate phenotypic data, 21,575 SNPs common to all the subpopulations were used. Phenotypic values were simulated as follows:

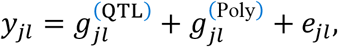

where *y*_*jl*_ corresponds to the phenotypic value of accession *l* from subpopulation *j*, 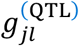 is the QTL contribution to the genotypic value, 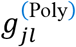 is the polygenic contribution to the genotypic value, and *e*_*jl*_ is an experimental error term, with *j =* 1,2,3 for China, Japan, and Korea subpopulations, respectively.

The polygenic effect 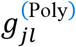 can be decomposed into two components as follows.

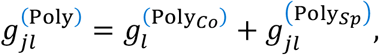

where 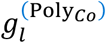 and 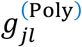 are respectively defined as the common and population specific polygenic components, with the following expressions:

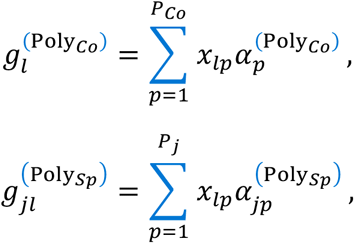

Here, 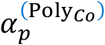 and 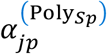 correspond to the *p*^th^ polygenic marker effect common to all population or specific to the *j*^th^ population, respectively. These effects were randomly sampled from 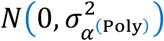. *P*_*Co*_ and *P*_*j*_ are the numbers of polygenes for the corresponding polygene types. Also, *x*_*lp*._ ∈ {−1,0,1} is the genotype score of accession *l* at marker *p*.

Similarly, 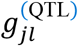 breaks down into three components as follows:

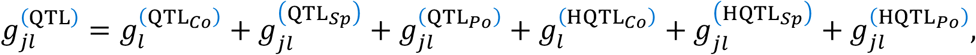

where 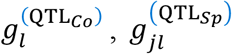 are defined as the sums of QTL effects common to all populations (hereinafter, referred to as Common QTLs) and specific 210 to the *j*^th^ population, respectively:

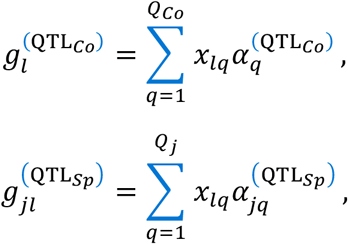

Here, 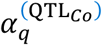 and 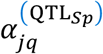 are the effects of *q*^th^ Common and *j*^th^ population-specific QTLs. Also, *Q*_*Co*_ and *Q*_*j*_ are the numbers of Common and specific QTLs. The population-specific QTLs account for population-specific interaction effects between QTLs and the discrete population structure. In this study, we only assumed the existence of China-specific QTLs (hereinafter, referred to as China QTLs) and no population-specific QTLs for Japan and Korea populations, i.e., *Q*_2_ *= Q*_3_ *=* 0. Finally, 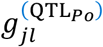 is defined as the sum of QTL effects proportional to the total polygenic effects 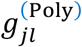 (hereinafter, referred to as Polygenes QTLs), as in the following equation.

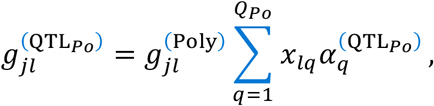

where 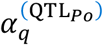 is a *q*^th^ Polygenes QTL effect, and *Q*_*Po*_ is the number of Polygenes QTLs. These Polygenes QTLs were introduced to express the interaction effects between QTLs and the continuous population structure.

We also considered haplotypic QTLs whose effects depend on the HBs they belong to. To simulate the haplotypic QTL effects, 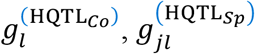, and 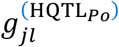, we first performed the “k-medoids” methods [53] for the markers in the HB of interest with *k =* 2 and divided the accessions into two haplotype groups. Then, we assume accessions in one haplotype group with the larger number of alternative alleles have some effects for that QTL, whereas those in the other group have no effects. These haplotypic QTLs were simulated for each QTL type of Common, China, and Polygenes, and are hereafter referred to as HB-Common, HB-China, and HB-Polygenes. An example of HB-China QTL is illustrated in S3 Fig.

We prepared four different scenarios depending on the simulation parameters, such as the number of QTLs (Table 1). As other parameters common to all the scenarios, we set the numbers of polygenes *P*_*Co*_ *=* 2,800 and *P*_*Ch*_ *= P*_*Ja*_ *= P*_*Ko*_ *=* 600. Also, each QTL effect was set as 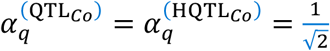 and 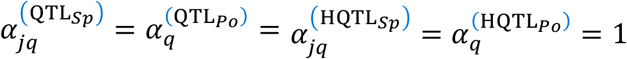, where 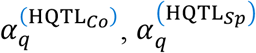, and 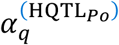 were *q*^th^ HB-Common, HB-China and HB-Polygenes QTL effects, respectively. Then, after the computation of the sum of QTL effects, the variance of polygenic effects 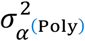 was automatically determined by setting the ratio between the QTL variance and polygenic variance (Var[**g**(^QTL^)]: Var[**g**(^Poly^)]) as in Table 1. Finally, the error term *e*_*jl*_ was randomly sampled from 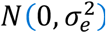, where the error variance 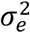 was automatically determined to set the heritability 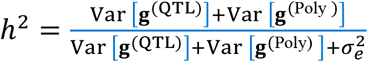 at either 0.75 or 0.8, depending on the scenario (Table 1).

**Table 1.** Simulation parameters for four different scenarios assumed in this study.

| Parameter Names | Scenario 1 | Scenario 2 | Scenario 3 | Scenario 4 |
| --- | --- | --- | --- | --- |
| Number of Common QTLs | 1 | 1 | 1 | 2 |
| Number of China QTLs | 1 | 1 | 0 | 0 |
| Number of Polygenes QTLs | 2 | 1 | 2 | 2 |
| Number of HB-Common QTLs | 1 | 1 | 1 | 0 |
| Number of HB-China QTLs | 1 | 1 | 0 | 0 |
| Number of HB-Polygenes QTLs | 2 | 1 | 2 | 0 |
| $\text{Var}[\mathbf{g}^{(\text{QTL})}]: \text{Var}[\mathbf{g}^{(\text{Poly})}]$ | 2:1 | 3:1 | 2:1 | 1:1 |
| Heritability | 0.75 | 0.8 | 0.75 | 0.8 |

For each scenario, the phenotypic data was simulated 100 times and used to evaluate the different GWAS model described in the **Comparison of five models** section.

### Proposed GWAS models

We introduce three SNPxGB GWAS models to cope with the presence of an unknown and possibly cryptic genetic structure in the data. Compared to the conventional SNP-based GWAS model, these SNPxGB models explicitly include an interaction term between a SNP and the genetic background. These models are then straightforwardly extended to account for HBxGB interactions.

#### 1. k-means (KM)

We first consider a classical SNPxGB interaction model. For a given marker *m*, one has

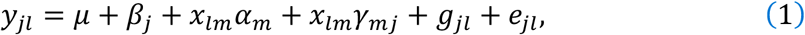

where *y*_*ji*_ is a phenotypic value of accession *l* from subpopulation *j, µ* is an intercept, *β*_*j*_ is a general effect of subpopulation *j, x*_*lm*_ ∈ {−1,0,1} is the genotype of accession *l* at marker *m, α*_*m*_ is the additive effect of allele 1 at marker *m*, and *γ*_*mj*_ is the additive population-specific effect of allele 1 in population *j* at marker *m, g*_*jl*_ is the polygenic effect of accession *l*, and 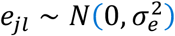 is an error term.

Here, *µ, β*_*j*_, *α*_*m*_, and *γ*_*mj*_ are fixed effects, and *g*_*jl*_ is a random effect that follows a multivariate normal distribution. One has

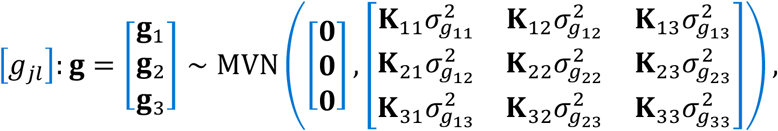

where **g**_*j*_ is the vector of polygenic effects of accessions from subpopulation *j*, **K**_*jj*′_ is an additive genomic relationship matrix between accessions from subpopulations *j* and *j*^′^ [51], which was estimated based on the marker genotype of 30,475 SNPs using the calcGRM function in the R package RAINBOWR version 0.1.31 [38], and 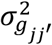 is the corresponding genetic variance/covariance.

As the genetic structure is assumed to be unknown, it must be inferred beforehand. Here we used the k-means algorithm to infer each cluster for the estimation of the general effect β_*j*_ or the population-specific effect γ_*mj*_ This k-means clustering was performed on the first 10 principal components of the principal component analysis (PCA), and each estimated cluster was regarded as a subpopulation. Here, we fixed the number of clusters at *k =* 3 which corresponds to the true number of subpopulations, i.e., we did not consider the problem of choosing *k* in this analysis.

In the SNPxGB model, after the estimation of marker effects, all the marker effects, including the interaction effect, are tested simultaneously, as in the null hypothesis of the following equation.

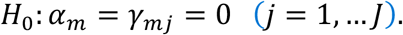

#### 2. Principal component analysis (PC)

As the second method, we accounted for the SNPxGB interaction by performing a PCA on the genomic relationship matrix **K** and using the first *B* principal component scores as interaction factors. The resulting model is

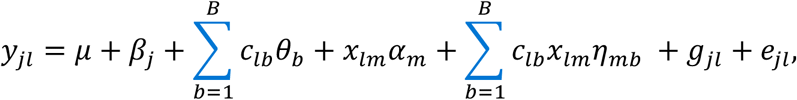

where *c*_*lb*_ is the *b*^th^ principal component score of accession *l, θ*_*b*_ is the covariate effect corresponding to the *b*^th^ score, and *η*_*mb*_ is the interaction effect between allele 1 of marker *m* and the *b*^th^ score. In this study, we only considered *B =* 2 for the PC model. Here, the null hypothesis is that the marker effect *α*_*m*_ and the interaction effects *η*_*mb*_ are equal to 0, i.e., *H*_0_: *α*_*m*_ *= η*_*mb*_ *=* 0.

#### 3. Discriminant analysis of principal components (DA)

As the third method, we utilized a discriminant analysis of principal components [52]. A PCA was performed on the genomic relationship matrix **K**, followed by a linear discriminant analysis on the retained first *P =* 10 principal components. The resulting model is

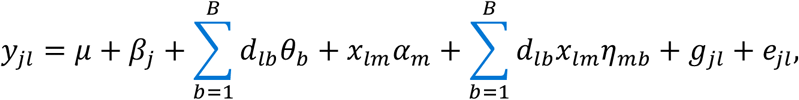

where *d*_*lb*_ is a *b*^th^ linear discriminant score of accession *l*. Here, since we used the estimated genetic background by the k-means clustering with *k =* 3 as the objective variable of the linear discriminant analysis, we obtained *B = k* − 1 linear discriminant scores. To perform the discriminant analysis of principal components, we used the dapc function in the R package adegenet version 2.1.4 [53,54].

### HBxGB model

The HBxGB model is the extension of the HB-based model [38] obtained by incorporating the interaction term between HB and a continuous population structure. Thus, the proposed HBxGB model explicitly considers the interaction term between an HB and polygenes *𝒱*_*lb*_ as in Eq 2.

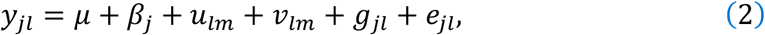

where *u*_*lb*_ and *𝒱*_*lm*_ are random effects for an HB of interest, and the other terms are described in Eq 1. *u*_*lb*_ is an *m*^th^ HB effect of accession *l* and is treated as a random effect as follows.

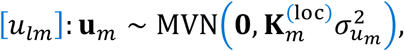

where 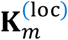 is a *L* × *L* genomic relationship matrix as for the *m*^th^ HB, and 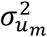 is a genetic variance of the *m*^th^ HB. In this study, as well as **K**, 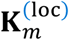 was estimated by the calcGRM function in the R package RAINBOWR version 0.1.31 [38]. However, since the estimation is based only on the marker genotype that belonged to the *m*^th^ HB, 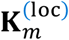 can be regarded as a local relationship matrix.

Next, *𝒱*_*lm*_ is an *m*^th^ HB effect interacting with the polygenes, which are assumed to follow a multivariate normal distribution as follows.

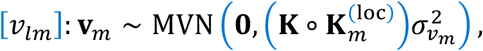

where 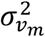 is a genetic variance of the *m*^th^ HB interacting with polygenes, and **A** ∘ **B** represents an Hadamard product between the two matrices **A** and **B**. Since **K** can be regarded as a global relationship matrix, the Hadamard product 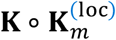 represents the interaction between the global and local genomic relationship matrices.

### Evaluation of the proposed models

#### Comparison of five models

In this study, we compared the following five GWAS models: the conventional SNP-based model (SNP) [21], the M2 model proposed by Rio et al. (M2) [27], the different SNPxGB models (KM, PC, and DA), the conventional HB-based GWAS model (HB) [38], and the HBxGB model (HBxGB). Since the SNPxGB and HBxGB models were already described above, we first briefly summarize the other three models as follows.

Here, the conventional SNP-based model used in this study was identical to the SNPxGB model in Eq 1, but with the interaction term omitted. Similarly, the HB-based GWAS model was the same as the HBxGB model in Eq 2 except for interaction term.

The M2 model proposed by Rio et al. [27] assumes that each marker effect depends not only on its SNP allele but also on its ancestry, i.e., country of origin/subpopulation in this study. Thus, the M2 model can be expressed as in the following equation.

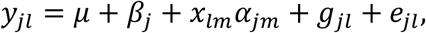

where *α*_*im*_ is the *m*^th^ marker effect from subpopulation *j*, and all the other terms are described in Eq 1. In this study, we assumed the M2 model could access the prior information on the subpopulations, as in [27], to evaluate the potential of the SNPxGB model without prior information. In the original article, several contrasts were considered for population-specific effects as in Eq 3, but also for common and divergent effects across subpopulations as in Eqs 4 – 7 for each marker:

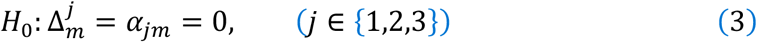

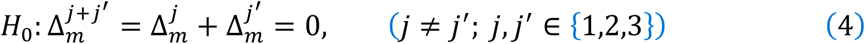

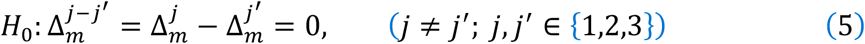

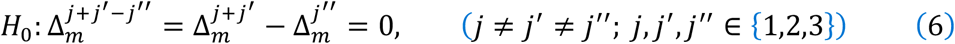

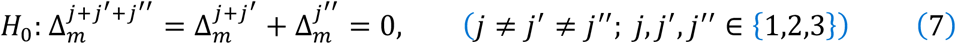

where there were three tests for Eqs 3 – 6 each and only one test for Eq 7, resulting in thirteen hypothetical tests in total for this M2 model. In this study, since we prepared for three QTL types of Common, China, and Polygenes and did not consider QTLs with divergent effects nor QTLs only common to two subpopulations in the simulations, among the above hypothetical tests, we only performed the four tests, i.e., three tests in Eq 3 (hereinafter, referred to as “M2C”, “M2J”, and “M2K” for *j =* 1,2,3, respectively) and one test in Eq 7 (hereinafter, referred to as “M2CJK”), to detect population-specific and Common QTLs, respectively. Also, we tested all the hypothetical tests in Eqs 3 – 7 simultaneously (hereinafter, referred to as “M2ALL”) to see whether the M2 model can detect Polygenes QTLs by any of the above tests.

#### Evaluation of the simulation results

Simulations were performed 100 times in order to evaluate for each GWAS model both its ability to control for false positive (FP) at the required nominal level and the detection performance. In a given simulation run, a marker/HB QTL was declared as detected if any marker belonging to the same LD block (as defined by PLINK1.9, see the Materials and marker genotype data subsection) had a significant *p*-value after controlling for multiple testing. In what follows, the Benjamini Hochberg procedure was considered to control the false discovery rate (FDR) at a nominal level of either 0.05 or 0.01.

#### Control of false positives

In each simulation run, QTLs were simulated throughout the genome except on one chromosome, called the *H*_0_ chromosome hereinafter. By utilizing the *H*_0_ chromosome, we first evaluated the FDR as the proportion of markers whose *p*-values were smaller than the significance level *α* among markers on the chromosome *H*_0_ in a given simulation run.

Additionally, as no QTL other than polygenes was simulated on chromosome *H*_0_, this chromosome was used to evaluate the FDR as the proportion of simulation runs in which a marker of the *H*_0_ chromosome was found to be significant.

#### Evaluation of detection power

Lastly, we computed a recall for each QTL type as follows to evaluate how each GWAS model could detect each QTL type.

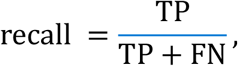

where FN is the number of false negatives and TP corresponds to the number of true QTLs which are not detected. These were defined at the LD block level in this study. In this study, we evaluated the recall for each QTL type, i.e., Common, China, Polygenes, HB-Common, HB-China, and HB-Polygenes, respectively.

#### Evaluation based on real phenotypes

We also validated the proposed models by applying them to real phenotypic data. This data, referring to two traits, i.e., oil and protein contents, came from the USDA Germplasm Resources Information Network database (www.ars-grin.gov). This information was organized by Bandillo et al. [40] and included phenotypes for a total of 12,116 *G. max* accessions. Out of these, 381 accessions, which were also part of the 900 accessions mentioned earlier, to evaluate the five GWAS models.

## Results

### Population structure of the soybean dataset

We used the soybean dataset from the USDA Soybean Germplasm Collection, which can be considered as a representative case of a panel that exhibits a cryptic and continuous population structure, as reported in Bandillo et al. [40]. Note that after selecting the 900 accessions, the analytical population for GWAS still had a continuous population structure, as shown in plots of the principal component analysis (Fig 1).

**Fig 1.**
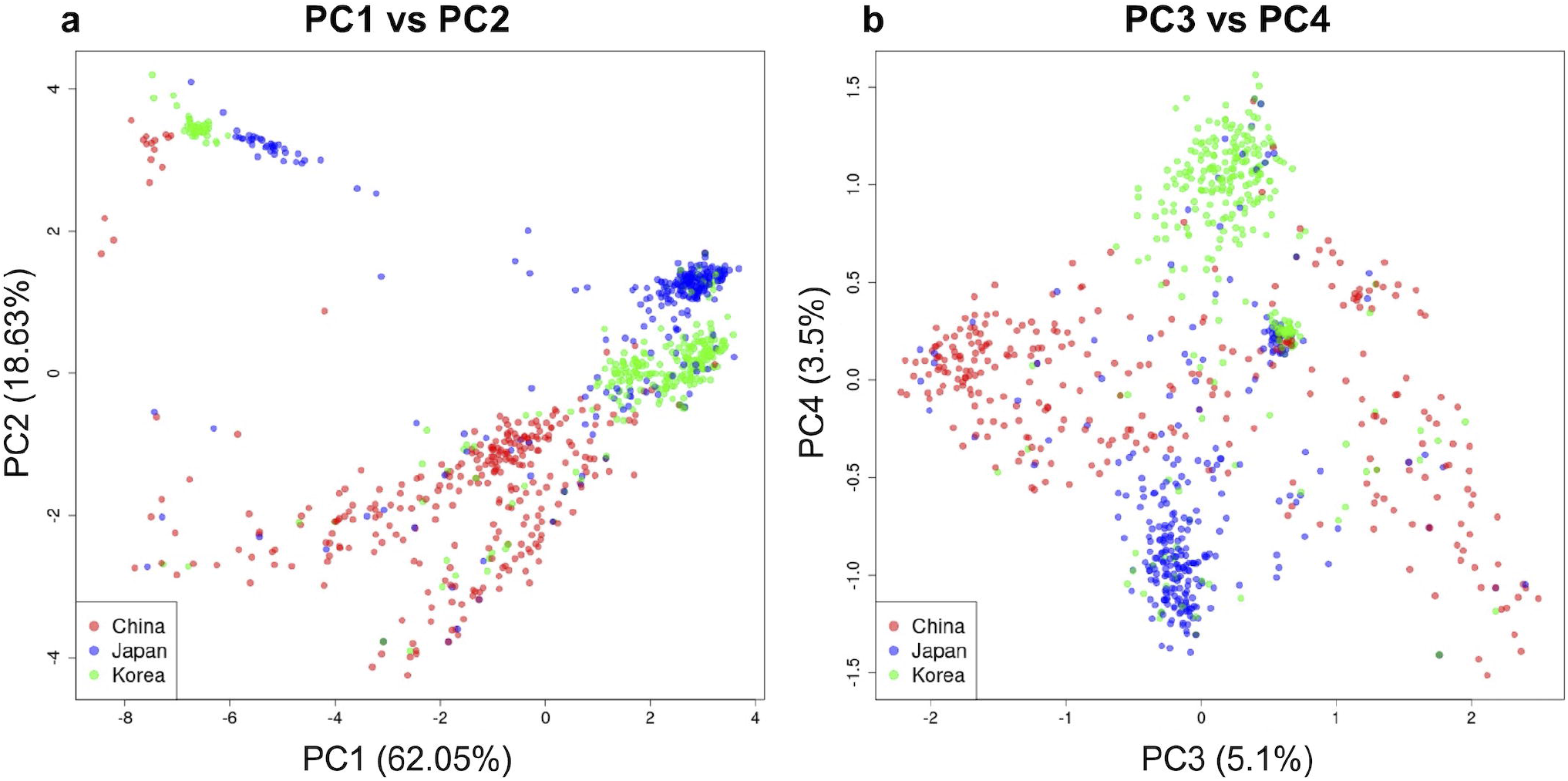
Principal Component Analysis Plot for the Analytical Population of the Selected 900 Accessions. Scatter plots of principal components PC1 vs PC2 (a) and PC3 vs PC4 (b). The red, blue, and green points correspond to the accessions from China, Japan, and Korea, respectively.

### Control of false positives

First, we plotted a boxplot for the first statistic, i.e., the proportion of markers whose *p* -values were smaller than the significance level *α* among markers on the chromosome *H*_0_ (Fig 2 for *α =* 0.05 and S4 Fig for *α =* 0.01). As a result, all the SNP-based GWAS models showed the value around the significance level *α*, whereas the HB-based GWAS models, HB and HBxGB, showed much smaller values than the significance level *α* in all the scenarios, especially for *α =* 0.05 (Fig 2). Thus, we can say that all the models could control the false positives, and especially the *p*-values of the HB-based models were deflated.

**Fig 2.**
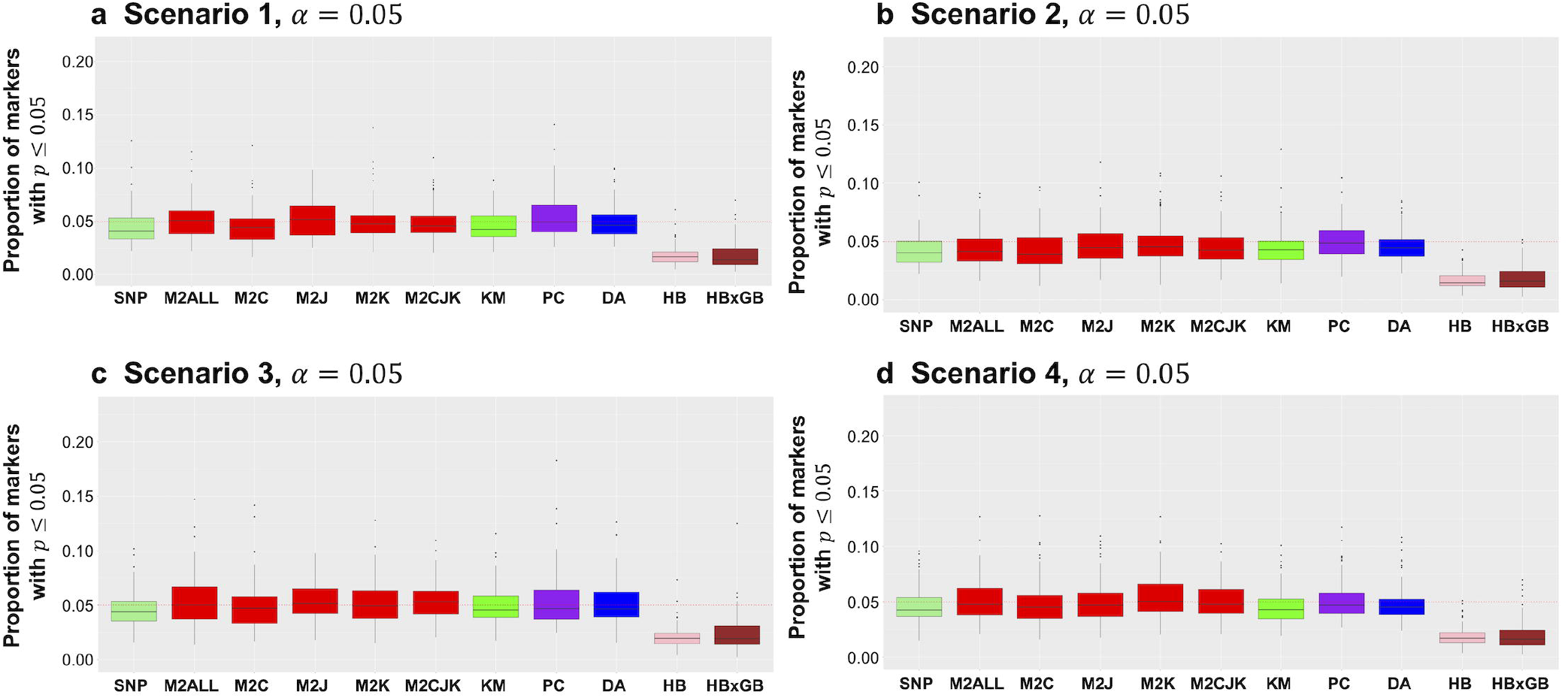
Proportion of markers with *p*-values lower than the significance level *α =* 0.05. (a) Scenario 1. (b) Scenario 2. (c) Scenario 3. (d) Scenario 4. The red dashed horizontal line correspond to the value of 0.05. The abbreviation of each model is as follows. SNP: The conventional SNP-based model [21]. M2ALL: The M2 model [27] testing for all the effects simultaneously. M2C: The M2 model [27] testing for the China-specific effects in Eq 3. M2CJK: The M2 model [27] testing for the Common effects in Eq 7. KM: The SNPxGB model with k-means. PC: The SNPxGB model with PCA. DA: The SNPxGB model with DAPC. HB: The HB model [38]. HBxGB: The HBxGB model.

Next, when we evaluated the mean of the FDRs based on the chromosome *H*_0_, all the models except HB in Scenario 3 for *α =* 0.01 showed lower FDR means than the significance level of 0.05 or 0.01 (S5 Fig). Thus, we can say that almost all the models could control the false positives well from the view of the second statistic.

### Evaluation of recalls

Recall was evaluated for each QTL type in each scenario. Figs 3 - 5 display the results for Common/HB-Common, China/HB-China, and Polygenes/HB-Polygenes, respectively, in Scenario 2 (see also S6 – S8 Figs for scenarios 1, 3, and 4).

**Fig 3.**
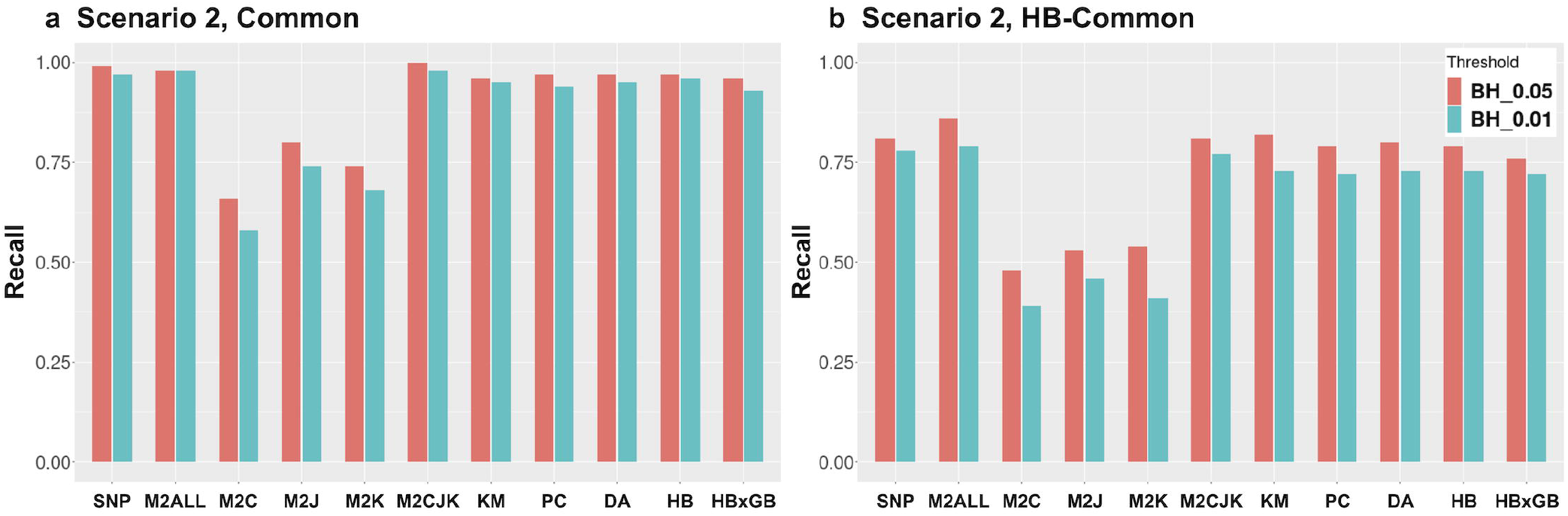
Recall for the Common and HB-Common QTLs in Scenario 2. (a) Common QTLs. (b) HB-Common QTLs. The red and blue bars show the results of the recalls after applying a Benjamini-Hochberg procedure to control Type I error rate at levels *α =* 0.05 and *α =* 0.01, respectively. The abbreviations of the models are the same as those of Fig 2.

First, all the tests except the M2 models testing for population-specific effects (M2C, M2J, M2K) succeeded in detecting the Common and HB-Common QTLs (Fig 3 and S6 Fig). The recall was generally lower for the HB-Common QTLs compared to the Common QTLs (Fig 3). Although the difference between the models was small, the M2 model testing all effects or common effects, the conventional SNP-based models, and the SNPxGB models generally showed almost the same recall, followed by the HB model, and the HBxGB model showed the lowest recall in Scenarios 1 and 2 (Fig 3 and S6 Fig a,b). On the other hand, in Scenarios 3 and 4, the HBxGB model showed almost the same recall as the SNPxGB models (S6 Fig c-e). Although the difference in recalls was slight, the best model differed between the scenarios among the SNP-based models. Also, the comparison between scenarios showed that the QTLs with larger variances, i.e., Scenarios 2 and 4 (Fig 3 and S6 Fig e), tended to show higher recall than those with smaller variances, i.e., Scenarios 1 and 3 (S6 Fig a-c), which was more pronounced for the HB-Common QTLs.

Next, the contrast between models was observed for the China and HB-China QTLs (Fig 4 and S7 Fig). The recall was generally lower for the HB-China QTLs compared to the China QTLs (Fig 4 and S7 Fig). For both QTL types assuming the interaction with discrete structure, the M2 model testing for the China-specific effects showed the highest recall, followed by the SNPxGB models, the HBxGB model, and then the conventional SNP and HB model showed the lowest recall (Fig 4). The recall for the QTLs with larger variances, i.e., Scenario 2 (Fig 4), was also larger than those with smaller variances, i.e., Scenario 1 (S7 Fig), for the China and HB-China QTLs.

**Fig 4.**
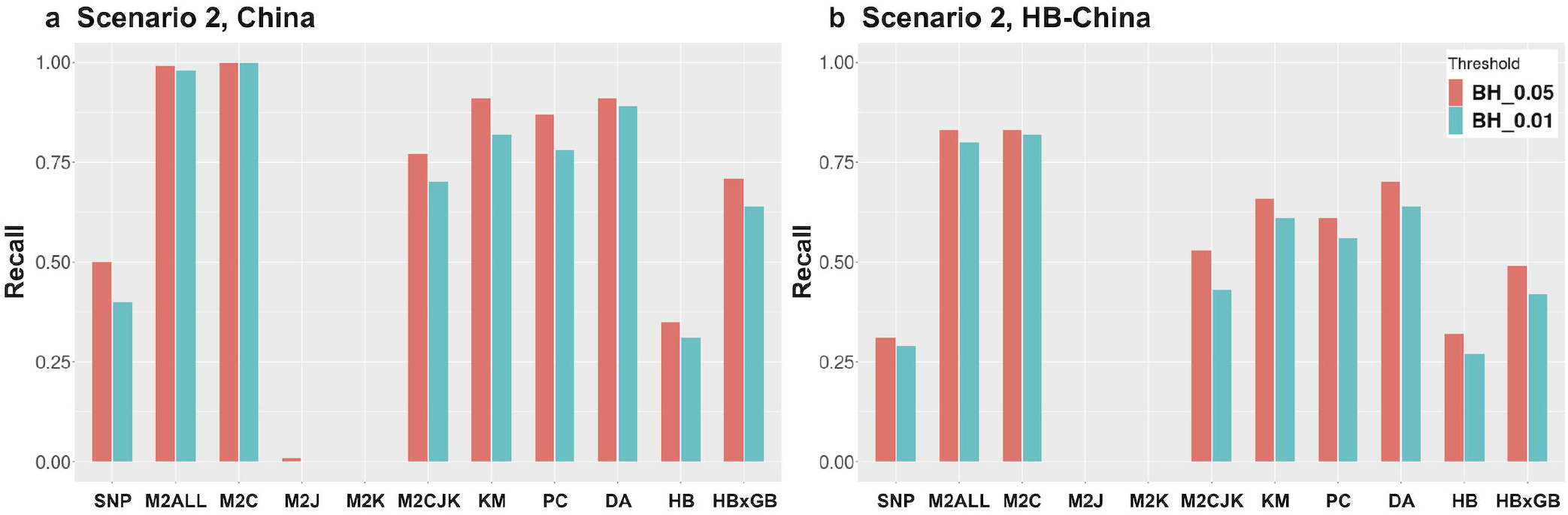
Recall for the China and HB-China QTLs in Scenario 2. (a) China QTLs. (b) HB-China QTLs. The red and blue bars show the results of the recalls after applying a Benjamini-Hochberg procedure to control Type I error rate at levels *α =* 0.05 and *α =* 0.01, respectively. The abbreviations of the models are the same as those of Fig 2.

Finally, the detection power significantly decreased in all the models for the Polygenes and HB-Polygenes QTLs, which assumed the interaction effects with a more continuous structure than the China and HB-China QTLs (Fig 5 and S8 Fig). The recall was generally lower for the HB-Polygenes QTLs compared to the Polygenes QTLs, particularly in the HBxGB model (Fig 5 and S8 Fig). Still, for both QTL types assuming the interaction with continuous structure, the proposed HBxGB model showed the highest recall, followed by the M2 model testing all the effects, the SNPxGB models (PC, DA, KM), and the HB and SNP models (Fig 5). Significantly, the HB and SNP models did not detect the Polygenes QTLs at all. The comparison between scenarios showed the superiority of the HBxGB model was more pronounced in the order of Scenarios 2 (Fig 5), 4 (S8 Fig e), 3 (S8 Fig c,d), and 1 (S8 Fig a,b), i.e., the order in which the greater variances were assigned to the Polygenes QTLs.

**Fig 5.**
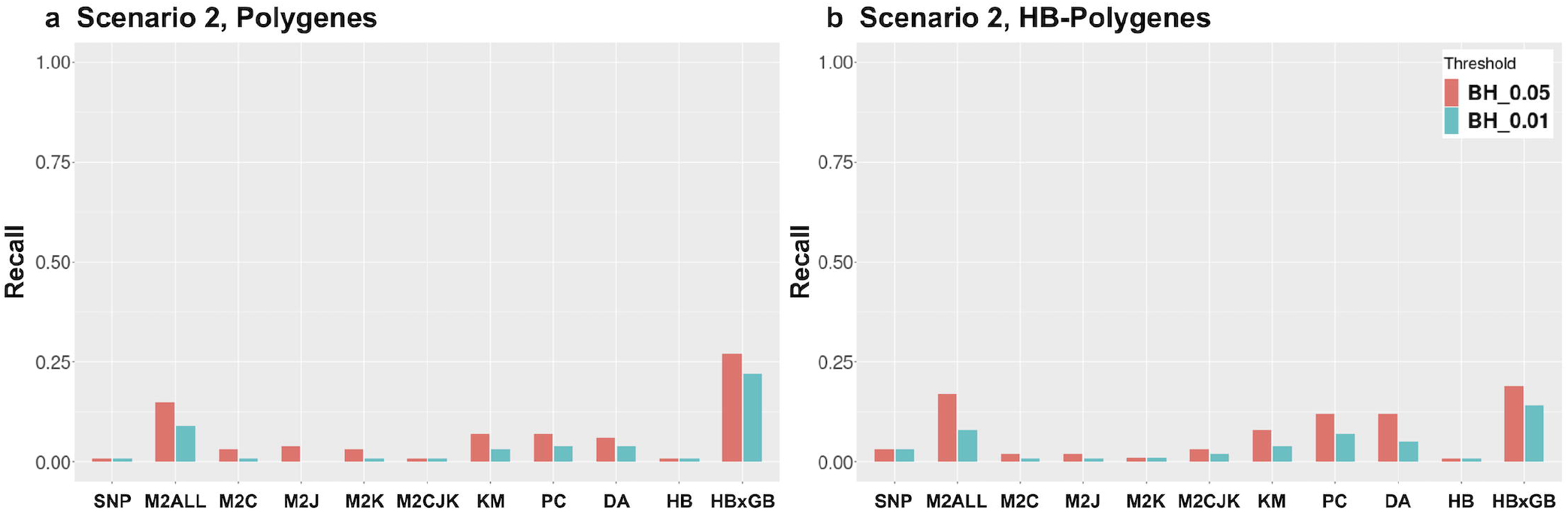
Recall for the Polygenes and HB-Polygenes QTLs in Scenario 2. (a) Polygenes QTLs. (b) HB-Polygenes QTLs. The red and blue bars show the results of the recalls after applying a Benjamini-Hochberg procedure to control Type I error rate at levels *α =* 0.05 and *α =* 0.01, respectively. The abbreviations of the models are the same as those of Fig 2.

### Application to real phenotypes

For each GWAS model, two levels of FDR, *α =* 0.05 and *α =* 0.10, were used to define each SNP as significantly associated. The numbers of SNPs/HBs and LD blocks detected by each GWAS model were summarized in S1 Table. Also, the numbers of SNPs/HBs and LD blocks uniquely detected by each GWAS model were summarized in S2 Table. For the oil content, the M2 model testing all the effects simultaneously (M2ALL) detected the most QTLs, followed by DA, M2CJK, SNP, HBxGB, and the other models detected no QTL (S1 Table). When focusing on the number of SNPs/HBs uniquely detected by each GWAS model, there were some SNPs/HBs that could be detected by only one method, i.e., M2ALL, DA, M2CJK, or SNP model (S2 Table). This result suggested that the GWAS models, including the proposed SNPxGB models, were complementary to detect QTL candidates for the oil content data. On the other hand, for the protein content, only the PC model detected markers associated with the protein (S1 and S2 Tables). For both traits, the number of detected SNPs/HBs increased when the FDR level was *α =* 0.10 compared to *α =* 0.05, especially for the SNP-based models.

Next, we plotted boxplots of the phenotypes for different alleles in each subpopulation (S9 Fig). Here, we showed one maker detected by multiple GWAS models for the oil (S9 Fig a), three markers detected by only one SNPxGB method (DA) for the oil (S9 Fig b-d), and two markers detected by only one SNPxGB method (PC) for the protein (S9 Fig e,f). While the marker detected multiple GWAS models for the oil seemed to have constant effects across subpopulations (S9 Fig a), the SNPxGB models could detect population-specific SNPs whose effects depended on genetic background, especially for the results of the oil content (S9 Fig b - d).

## Discussion

Population stratification is one of the most severe problems in GWAS, and it is known that QTL effects interacting with the population structure also affect the GWAS results. Although the model that accounted for such population-specific QTLs has been proposed so far [27], the model required access to the information on population structure and did not assure the detection of the QTLs whose effects depend on the continuous population structure. In this paper, we propose a novel GWAS model that can detect the causal genes interacting with the discrete and continuous population structure without any prior information on the population structure.

### Control of false postives in each GWAS model

First, when we focused on the two statistics computed based on chromosome *H*_0_ with the null hypothesis, i.e., the FDR and the proportion of markers with *p*-values less than the significance level *α*, almost all the models showed lower or close values to *α* in all the scenarios (Fig 2, S4 and S5 Figs). Since the FDR evaluated the control level of the false positives depending on whether the marker on the chromosome *H*_0_ was detected or not, from the above results, almost all the models could control the number of false positives well in terms of the FDR. Also, since the proportion of markers with *p*-values less than the significance level was used to check whether the distribution of *p*-values followed the uniform distribution under the null hypothesis, we could confirm that there was no overall inflation of the − log_10_(*p*) values for all the models in all the scenarios. On the other hand, the HB and HBxGB models showed much lower values than *α*, meaning that these haplotype-based models are more conservative than the SNP-based models. In summary, it was proved that all the models, including the proposed models, could control false positives from these two summary statistics.

### Detection power of each GWAS model

Next, when looking at the recall of each model, for the Common and HB-Common QTLs, the SNP-based models such as SNP, SNPxGB, and M2 generally showed a bit higher recalls than the HB-based models such as HB and HBxGB, especially in Scenarios 1 and 2 (Fig 3 and S6 Fig). For the China and HB-China QTLs, the M2 model showed the highest recall, followed by the SNPxGB models, HBxGB models, and then the other models without considering the interaction effects (Fig 4 and S7 Fig). Finally, the proposed HBxGB model was the most prominent in detecting these QTLs for the Polygenes and HB-Polygenes QTLs (Fig 5 and S8 Fig). From these results, it can be said that the models assuming the same situation as the simulation settings for each QTL type were suitable for detecting the corresponding QTLs.

### Characteristics of the SNPxGB models

When focusing on our first proposed models, the SNPxGB models, they showed higher recalls than the conventional SNP-based or HB-based models for the China and HB-China QTLs (Fig 4). Since the SNPxGB models aimed to detect effects interacting with the genetic background, these results for the China and HB-China QTLs were consistent with what we had expected. On the other hand, for these population-specific QTLs, the recall of the SNPxGB was lower than that of the M2 model testing the China effects. This occurred because the SNPxGB did not utilize the true information on population structure, while the M2 model did. It was clear that using the estimated clusters instead of true countries of origin affected the detection of these population-specific QTLs. In this study, the clusters were estimated by the k-means algorithm for simplicity, but future studies could use more sophisticated methods like Structure [55] or ADMIXTURE [56] analysis to narrow the gap between the true and estimated clusters and improve the detection power.

Among the three proposed SNPxGB models, the model with DAPC (DA) generally showed the highest recall for China and HB-China QTLs (Fig 4). Thus, it could be the best way to infer the population structure when we do not have prior information on the population structure. Also, the model with k-means (KM) showed higher recalls than that with PCA (PC), meaning that considering the population structure as a discrete one is more suitable for detecting QTLs interacting with the discretely distributed genetic background. On the contrary, for the Polygenes and HB-Polygenes QTLs, although the recalls of the SNPxGB models were not so high, we can confirm the trend that the SNPxGB model assuming the interaction with more continuous structure, i.e., PC and DA, showed higher recalls than KM (Fig 5). From these results, the SNPxGB models were more suitable for detecting population-specific QTLs, and the best model among these models differed for the different types of QTLs by reflecting the characteristics of each model, i.e., assumptions for the interaction with discrete or continuous population structure. In addition, although we only assumed additive QTLs interacting with discrete structure in the SNPxGB models, the M2 and M3 models by Rio et al. have already extended to those considering the dominant effects so that they could be applied to factorial and admixed hybrid populations [57]. Thus, extending our SNPxGB models to those considering non-additive effects can be one of our future tasks.

### Characteristics of the HBxGB model

Next, when focusing on our second HBxGB model, for the Common and HB-Common QTLs, the recall was almost the same as the HB model and a bit lower than the SNP-based models (Fig 3). This is because the HB-based models tended to return more conservative results than the SNP-based models, as seen in the results for the proportion of markers with *p*-values lower than the significance levels (S4 Fig). This seems to be a weakness of the HB-based models in view of the recall, but it also leads to their advantage in the aspect of controlling false positives [38]. For the China and HB-China QTLs, the HBxGB model showed lower recall than the models to detect the population-specific QTLs, i.e., the M2 model and the SNPxGB models, but higher recalls than the conventional SNP-based and HB models (Fig 4). This result suggested that although the proposed HBxGB model could capture the population-specific QTL effects at a certain level, it was not specialized in detecting the population-specific QTLs, and superior models to the HBxGB existed for those QTLs interacting with discrete structure. Finally, as seen above, for the Polygenes and HB-Polygenes QTLs, the developed HBxGB model showed the highest recall in all the scenarios (Fig 5). This result was consistent with our expectation that the proposed HBxGB model could detect QTLs interacting with continuous population structure. In other words, considering interaction effects by the Hadamard product between the local and global relationship matrices led to efficient detection of the epistasis between QTLs and polygenes. From these aspects, it can be said that we have successfully developed a new GWAS model that can detect the Polygenes QTLs, which have been challenging to detect with previous SNP-based models, including the M2 model that can use the prior information on the population structure.

### Detection of haplotypic QTLs

When looking at the haplotypic QTLs, the recalls for the haplotypic QTLs were generally lower than those for the normal QTLs with the same QTL type, i.e., the recalls for the HB-Common, HB-China, and HB-Polygenes QTLs (Figs 3b, 4b, and 5b) were lower than those for the Common, China, and Polygenes QTLs, respectively (Figs 3a, 4a, and 5). This decrease in recalls for the haplotypic QTLs can be attributed to the reduction in allele frequency compared to the normal QTLs [58]. This problem is closely linked to the difficulty in detecting rare alleles or variants, i.e., alleles with low frequencies, which play an essential role in complex traits and have been revealed to have more significant genetic effects than common variants [59], not only in humans [60,61] but also in plants [62–64]. Since these rare alleles are hard to detect from a statistical point of view, many models have been developed to detect such significant variants using the SNP-set/haplotype-based approach [65–69], including the conventional HB model [38]. Thus, we had expected the HB-based models, such as the HB and HBxGB, would benefit from the detection of the haplotypic QTLs compared to the other SNP-based models, but from the results in this study, the comparison between the models for the normal and haplotypic QTLs showed almost the same trend, and we could not confirm the superiority of the HB-based models over the SNP-based models for the haplotypic QTLs. In particular, when comparing the HB-Polygenes QTLs with the Polygenes QTLs, the recall of the HBxGB model decreased the most among all the models. These results suggested that the HB-based models were not necessarily suitable for detecting haplotypic QTLs assumed in this study. Thus, we should develop a new GWAS approach for future tasks that can efficiently detect the QTLs whose effects depend on the haplotype where they belong. In order to achieve this new GWAS model for detecting the haplotypic QTLs, not only the interaction between QTLs and population structure but also the interaction between QTLs and haplotype should be explicitly included in the model in the future.

### The comparison between different scenarios

Here, we discuss the comparison across the results in different scenarios. First, for the Common and HB-Common QTLs, the HBxGB model showed relatively higher recalls in the comparison across models in Scenarios 3 and 4 than in the other two scenarios (Fig 3 and S6 Fig). This result was deeply related to the existence of the China and HB-China QTLs. With the presence of the population-specific QTLs, such as China and HB-China QTLs, as in Scenarios 1 and 2, the Common QTLs were confounded with the population-specific QTLs by the HBxGB model, which resulted in lower recalls than without the population-specific QTLs as in Scenarios 3 and 4. This result was caused by the characteristics of the HBxGB that did not separate the common and population-specific effects explicitly in the model, and thus, by combining the idea of the SNPxGB models, the HBxGB model is expected to be extended to the model that can capture not only the common but also the QTL effects interacting with discrete and continuous structure in the future. Next, for all the QTL types in all the models, the larger the genetic variance accounted for by the QTL, the higher the recall (Figs 3 – 5, S6 – S8 Figs). In particular, this trend differed across models for the Polygenes QTLs (Fig 5a), and we can see the HBxGB model was indeed able to detect the Polygenes QTLs compared to the other models in Scenario 2 with the large QTL variance. These results in different scenarios also helped us understand the characteristics of our proposed models.

### Application to real phenotypes

Finally, when focusing on the GWAS results of the real phenotypes for the oil and protein contents, some GWAS models could detect SNPs/HBs (S1 Table). Although the M2 model testing all the effects simultaneously (M2ALL) detected the most QTLs for the oil content (S1 Table), other GWAS models, including one of the SNPxGB models, DA, detected SNPs that could not be detected by the M2ALL (S2 Table). This result suggested that the GWAS models, including the proposed models, complementarily contributed to detecting QTL candidates for the real phenotypes. Also, from the boxplots of the phenotypes for different alleles, we can see that the SNPxGB models could detect some markers whose effects differed between subpopulations (S9 Fig b - d). However, since these markers could not be detected by other models, such as M2ALL, they may not be simple population-specific QTLs that can be detected by the previous GWAS models.

When focusing on the marker detected by the HBxGB model for the oil content, this marker was also detected by other models, i.e., the conventional SNP-based and the M2 model testing for the common effects (S9 Fig a). Thus, this marker can be said to have common effects, which can also be confirmed from the boxplot (S9 Fig a). However, since this marker could not be detected by the HB model with a threshold at FDR of 0.05, the relative detection power of the HBxGB seemed to be larger than that of the HB model. The presence of this marker does not necessarily directly indicate the usefulness of HBxGB, but it suggests the presence of QTLs that indicate slight interaction with some continuous population structure, in addition to common effects. From these results, although we cannot say that the HBxGB is always the best model, we should try various GWAS models, including the proposed SNPxGB and HBxGB models, since these models are complementary in detecting QTLs associated with actual agronomical traits.

## Conclusion

In this study, we develop novel GWAS models, SNPxGB and HBxGB, to detect QTLs interacting with the discrete and continuous population structure without its prior information, respectively. The HBxGB model remarkably succeeded in detecting the QTLs interacting with the continuous structure, which could not be easily detected by the previous models developed to detect population-specific QTLs with the prior population structure information. Also, the results applied to the real phenotypes showed that the GWAS models complementarily contributed to the detection of QTL candidates, suggesting that we should try various types of GWAS models, including the SNPxGB and HBxGB models. By applying our proposed models to various types of actual data, the proposed model is expected to help us reveal the unknown genetic mechanism/architecture of many complex traits, which have not been realized with the previous GWAS models.

## Acknowledgments

This study was conducted during the study in France with the support of the JSPS Overseas Challenge Program for Young Researchers.

## Declarations Data availability

The genome sequencing data obtained with the Illumina Infinium SoySNP50K BeadChip for all genotypes are available at the website of “SoyBase” project (https://soybase.org/snps/index.php). The selected and simulated datasets in the present study are available from the “KosukeHamazaki/DLIPG” repository on GitHub, https://github.com/KosukeHamazaki/DLIPG.

## Code availability

The proposed SNPxGB and HBxGB models were implemented in the existing R package “RAINBOWR” from version 0.1.29. A stable version of RAINBOWR is available from the CRAN (Comprehensive R Archive Network), https://cran.r-project.org/web/packages/RAINBOWR/index.html. The latest version of RAINBOWR is also available from the “KosukeHamazaki/RAINBOWR” repository on GitHub, https://github.com/KosukeHamazaki/RAINBOWR. The scripts used in this study are available from the “KosukeHamazaki/DLIPG” repository on GitHub, https://github.com/KosukeHamazaki/DLIPG.

## Supporting information

**S1 Fig. Example of the population-specific QTL effects**.

**S2 Fig. Image of the conventional and developed GWAS models used in this study**. (a) The conventional SNP-based model. (b) The conventional HB-based model. (c) The developed SNPxGB model. (d) The developed HBxGB model.

**S3 Fig. Image of an example of the HB-China QTLs**.

**S4 Fig. Proportion of markers with *p*-values lower than the significance level** *α =* 0.01. (a) Scenario 1. (b) Scenario 2. (c) Scenario 3. (d) Scenario 4. The red dashed horizontal line correspond to the value of 0.01.

**S5 Fig. FDRs based on the chromosome under the null hypothesis**. (a) Scenario 1. (b) Scenario 2. (c) Scenario 3. (d) Scenario 4. The red and blue bars show the results of the FDRs for the Benjamini-Hochberg thresholds with the significance levels *α =* 0.05 and *α =* 0.01, respectively. The red and blue dashed horizontal lines correspond to FDR *=* 0.05 and FDR *=* 0.01, respectively. The abbreviations of the models are the same as those of Fig 2.

**S6 Fig. Recall for the Common and HB-Common QTLs in Scenarios 1, 3, and 4**. (a), (c), (e) Common QTLs. (b), (d) HB-Common QTLs. (a), (b) Scenario 1. (c), (d) Scenario 3. (e) Scenario 4. The red and blue bars show the results of the recalls after applying a Benjamini-Hochberg procedure to control Type I error rate at levels *α =* 0.05 and *α =* 0.01, respectively. The abbreviations of the models are the same as those of Fig 2.

**S7 Fig. Recall for the China and HB-China QTLs in Scenario 1**. (a) China QTLs. (b) HB-China QTLs. The red and blue bars show the results of the recalls after applying a Benjamini-Hochberg procedure to control Type I error rate at levels *α =* 0.05 and *α =* 0.01, respectively. The abbreviations of the models are the same as those of Fig 2.

**S8 Fig. Recall for the Polygenes and HB-Polygenes QTLs in Scenarios 1, 3, and 4**. (a), (c), (e) Polygenes QTLs. (b), (d) HB-Polygenes QTLs. (a), (b) Scenario 1. (c), (d) Scenario 3. (e) Scenario 4. The red and blue bars show the results of the recalls after applying a Benjamini-Hochberg procedure to control Type I error rate at levels *α =* 0.05 and *α =* 0.01, respectively. The abbreviations of the models are the same as those of Fig 2.

**S9 Fig. Boxplots of phenotypes of the oil and protein contents for the different alleles of the six detected markers**. (a) Marker detected by the multiple GWAS models for the oil content. (b), (c), (d) Markers uniquely detected by the DA model for the oil content. (e), (f) Markers uniquely detected by the PC model for the oil content. The title displays the marker name, followed by the names of models capable of detecting that marker. Models without parentheses detected the marker at an FDR threshold level of 0.05, while those with parentheses detected the marker at an FDR threshold level of 0.1.The abbreviations of the models are the same as those of Fig 2.

**S1 Table.**
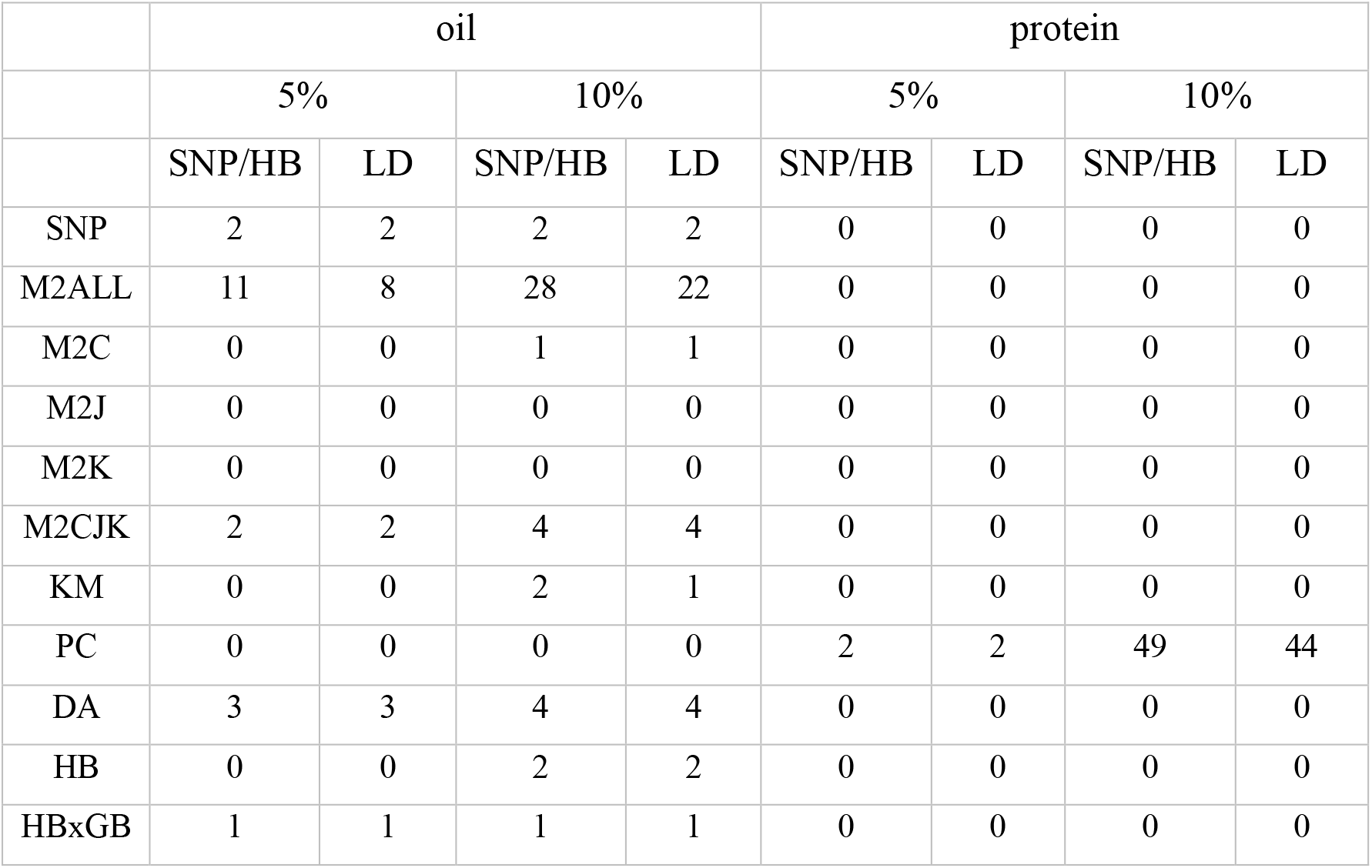
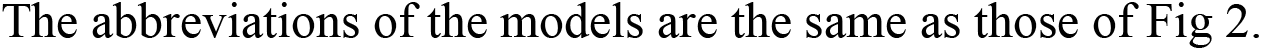
Numbers of SNPs/HBs and LD blocks detected by each GWAS model, using a FDR of 5% and 10%.

|  | oil |  |  |  | protein |  |  |  |
| --- | --- | --- | --- | --- | --- | --- | --- | --- |
|  | 5% |  | 10% |  | 5% |  | 10% |  |
|  | SNP/HB | LD | SNP/HB | LD | SNP/HB | LD | SNP/HB | LD |
| SNP | 2 | 2 | 2 | 2 | 0 | 0 | 0 | 0 |
| M2ALL | 11 | 8 | 28 | 22 | 0 | 0 | 0 | 0 |
| M2C | 0 | 0 | 1 | 1 | 0 | 0 | 0 | 0 |
| M2J | 0 | 0 | 0 | 0 | 0 | 0 | 0 | 0 |
| M2K | 0 | 0 | 0 | 0 | 0 | 0 | 0 | 0 |
| M2CJK | 2 | 2 | 4 | 4 | 0 | 0 | 0 | 0 |
| KM | 0 | 0 | 2 | 1 | 0 | 0 | 0 | 0 |
| PC | 0 | 0 | 0 | 0 | 2 | 2 | 49 | 44 |
| DA | 3 | 3 | 4 | 4 | 0 | 0 | 0 | 0 |
| HB | 0 | 0 | 2 | 2 | 0 | 0 | 0 | 0 |
| HBxGB | 1 | 1 | 1 | 1 | 0 | 0 | 0 | 0 |

The abbreviations of the models are the same as those of Fig 2.

**S2 Table.** Numbers of SNPs/HBs and LD blocks uniquely detected by each GWAS model, using a FDR of 5% and 10%.

|  | oil |  |  |  | protein |  |  |  |
| --- | --- | --- | --- | --- | --- | --- | --- | --- |
|  | 5% |  | 10% |  | 5% |  | 10% |  |
|  | SNP/HB | LD | SNP/HB | LD | SNP/HB | LD | SNP/HB | LD |
| SNP | 1 | 1 | 0 | 0 | 0 | 0 | 0 | 0 |
| M2ALL | 10 | 7 | 23 | 18 | 0 | 0 | 0 | 0 |
| M2C | 0 | 0 | 0 | 0 | 0 | 0 | 0 | 0 |
| M2J | 0 | 0 | 0 | 0 | 0 | 0 | 0 | 0 |
| M2K | 0 | 0 | 0 | 0 | 0 | 0 | 0 | 0 |
| M2CJK | 0 | 0 | 0 | 0 | 0 | 0 | 0 | 0 |
| KM | 0 | 0 | 2 | 1 | 0 | 0 | 0 | 0 |
| PC | 0 | 0 | 0 | 0 | 2 | 2 | 49 | 44 |
| DA | 3 | 3 | 4 | 4 | 0 | 0 | 0 | 0 |
| HB | 0 | 0 | 0 | 0 | 0 | 0 | 0 | 0 |
| HBxGB | 0 | 0 | 0 | 0 | 0 | 0 | 0 | 0 |

The abbreviations of the models are the same as those of Fig 2.

## Author contributions

KH, HI, and TMH conceived and designed the study. KH and TMH developed the new models. KH prepared for the dataset, performed the simulation of phenotypes, conducted the mathematical and statistical analyses, and summarized the results with visualization. TMH supervised the study. KH and TMH wrote the manuscript in consultation with HI. All authors have read and approved the final manuscript.

## Competing interests

Not applicable. The authors declare that there are no conflicts of interest.

## Ethics declarations

Not applicable.

## Notes

https://github.com/KosukeHamazaki/DLIPG

### Competing Interest Statement

The authors have declared no competing interest.

