## Supplementary figures and images for "Development of a novel GWAS method to detect QTL effects interacting with the discrete and continuous population structure"

### S1 Fig

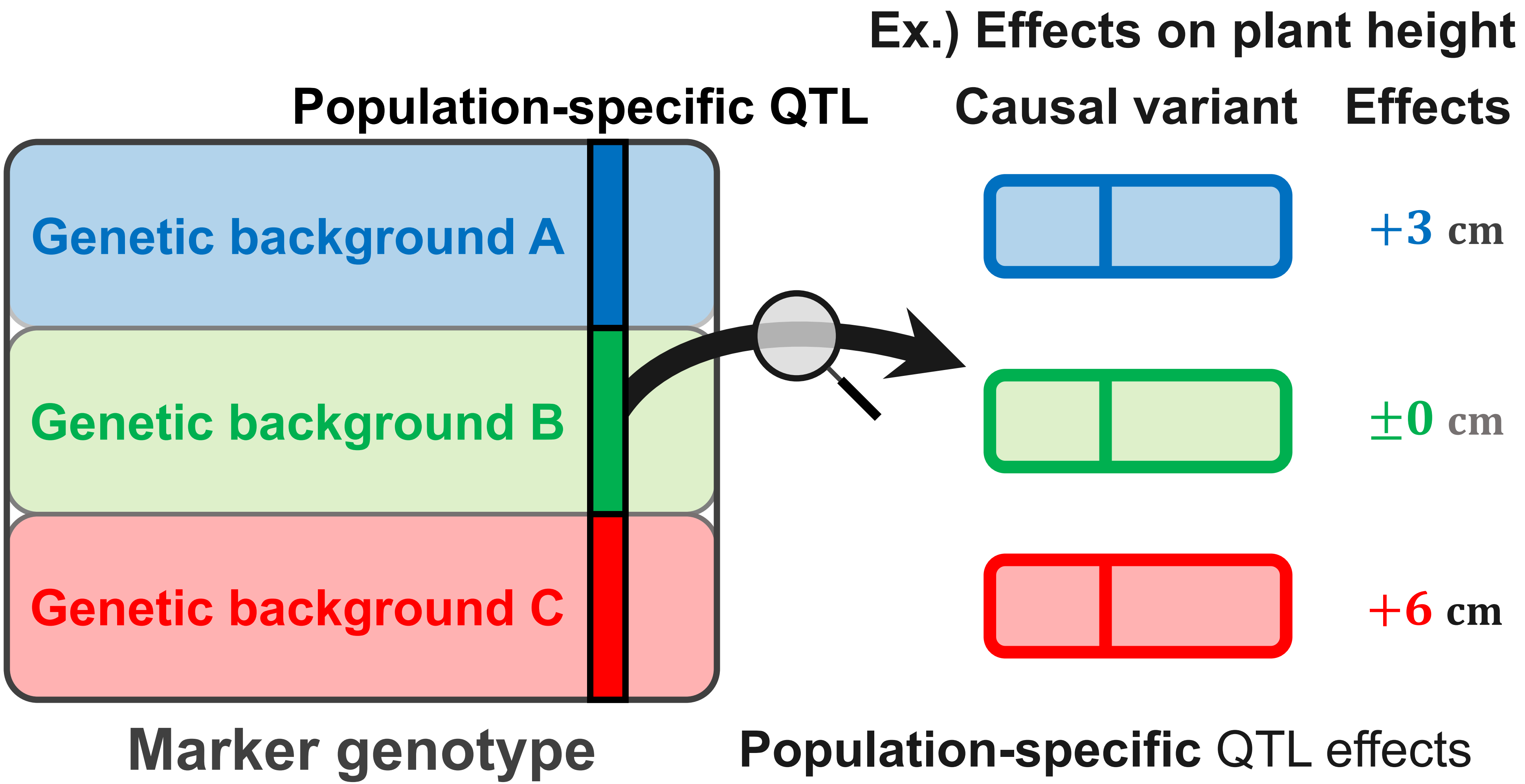

### S2 Fig

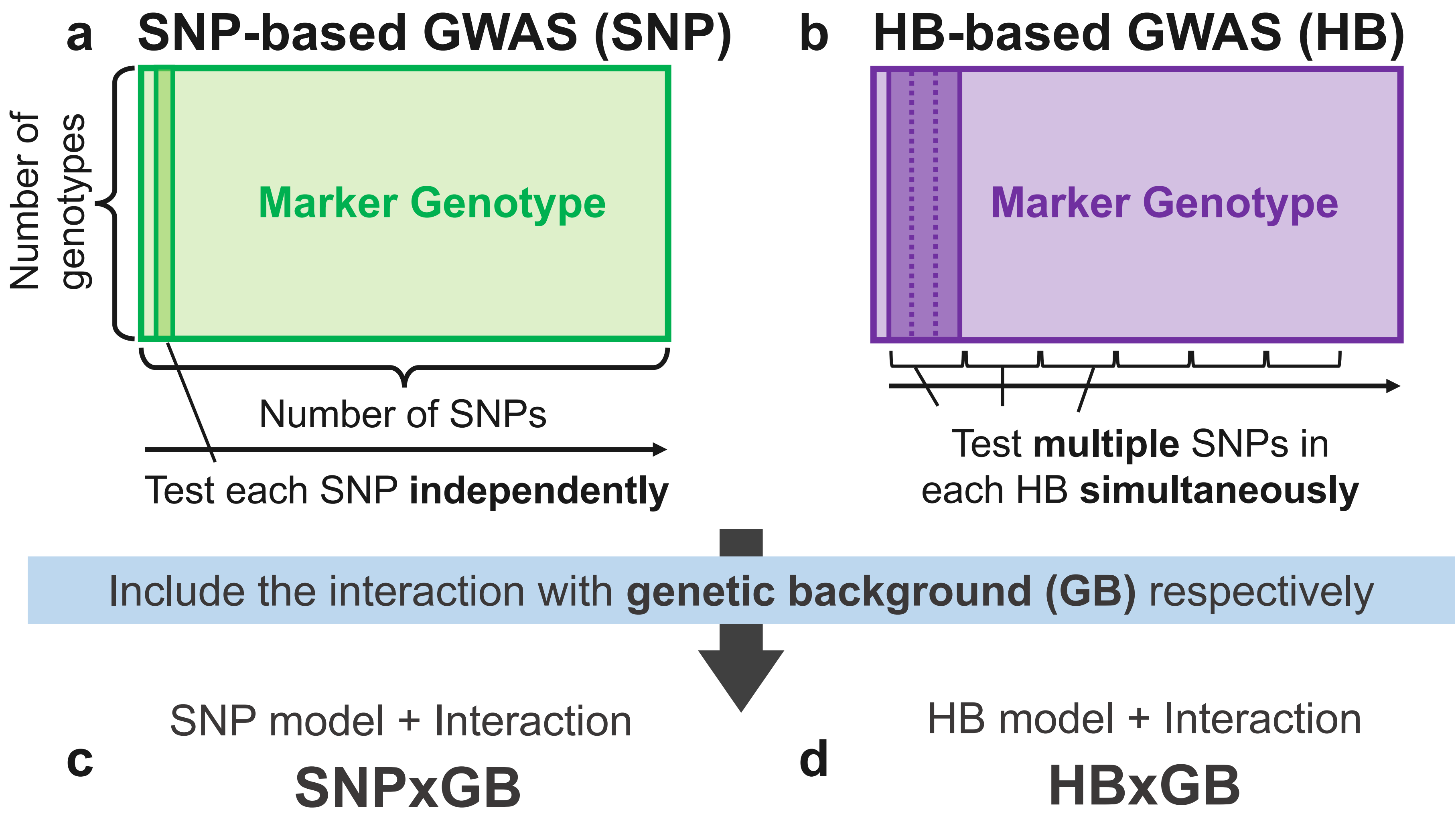

### S4 Fig

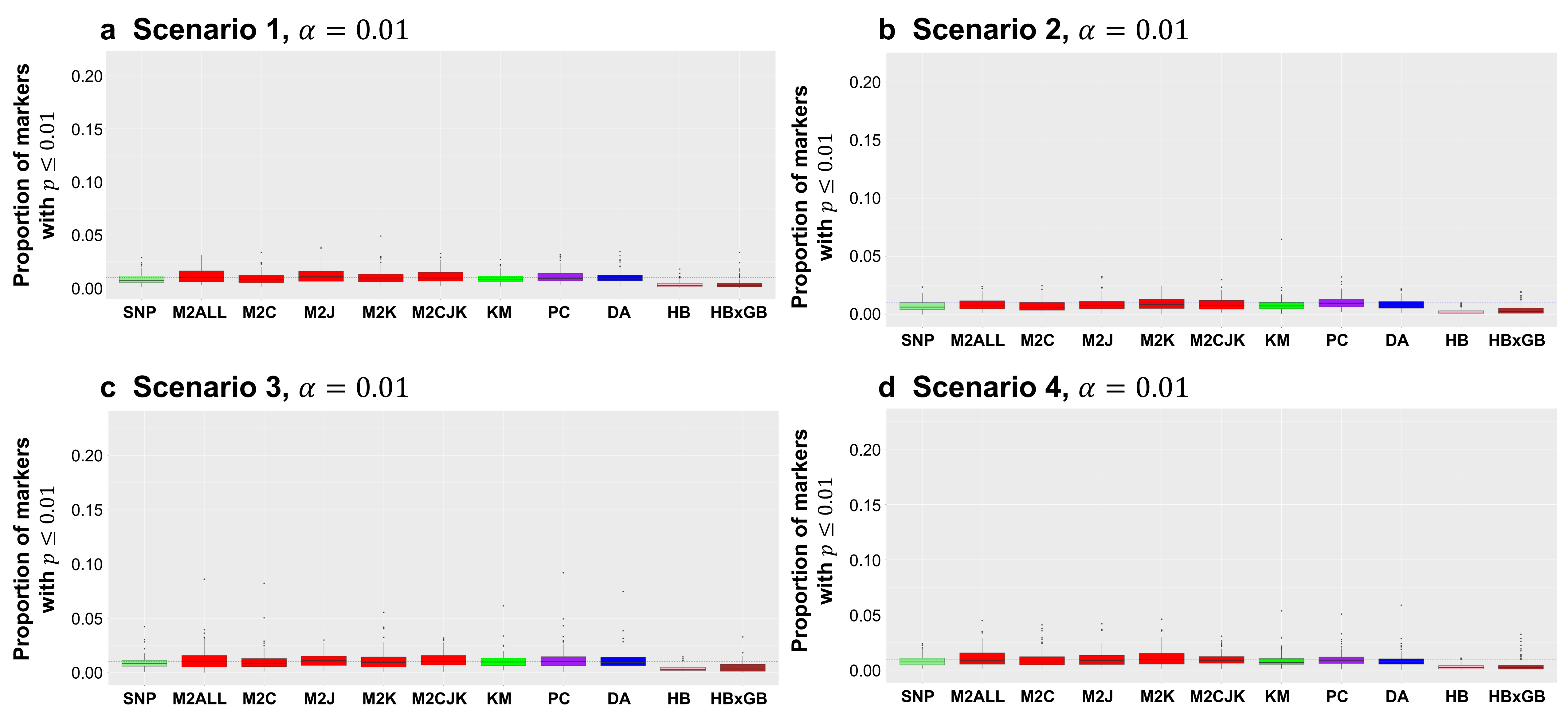

### S5 Fig

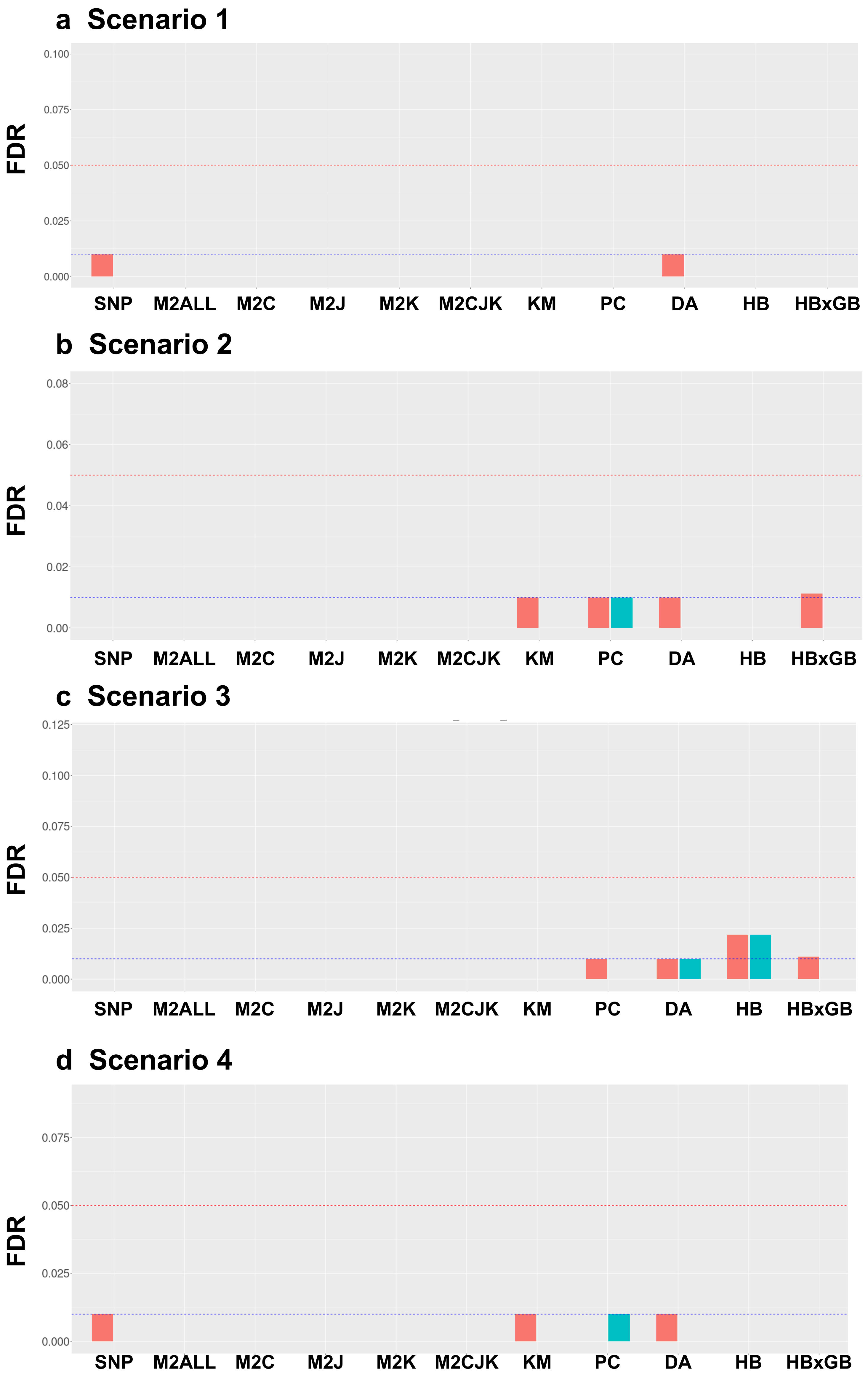

### S6 Fig

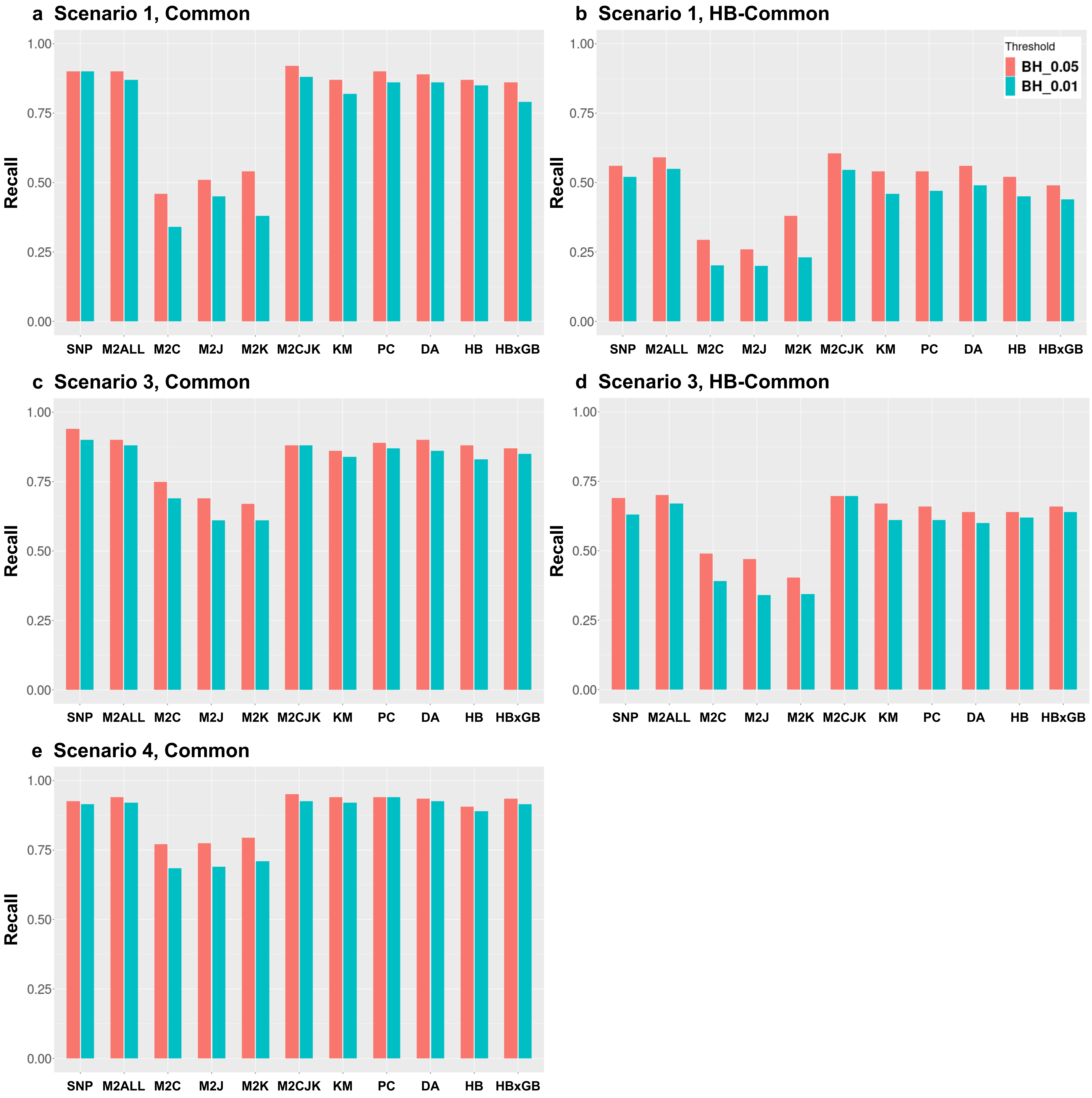

### S7 Fig

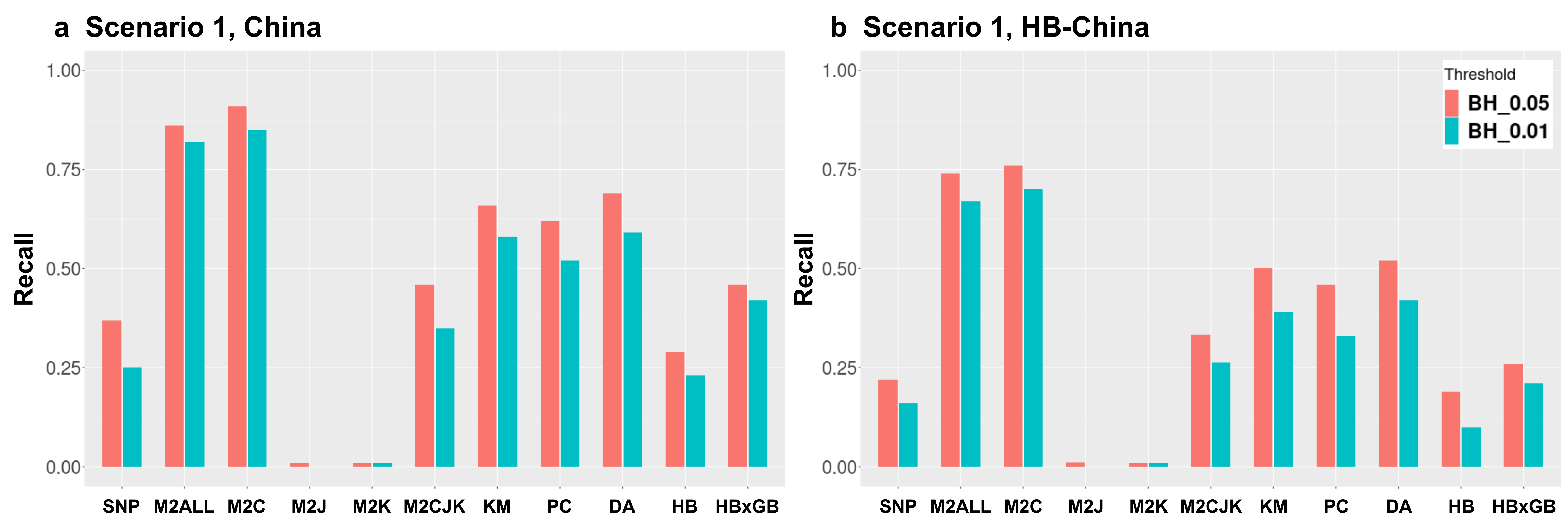

### S8 Fig

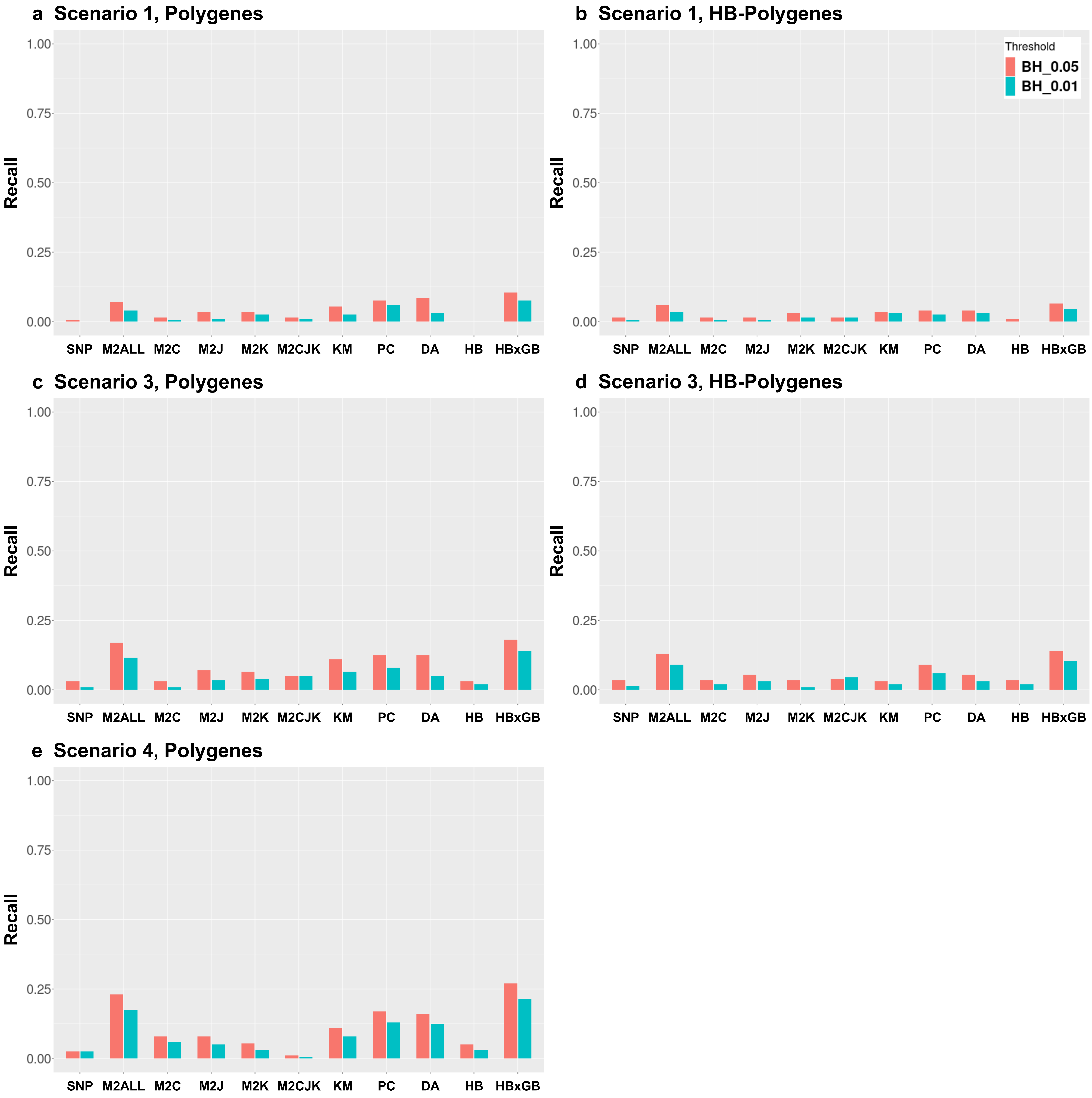

### S9 Fig

#### (1) Oil content

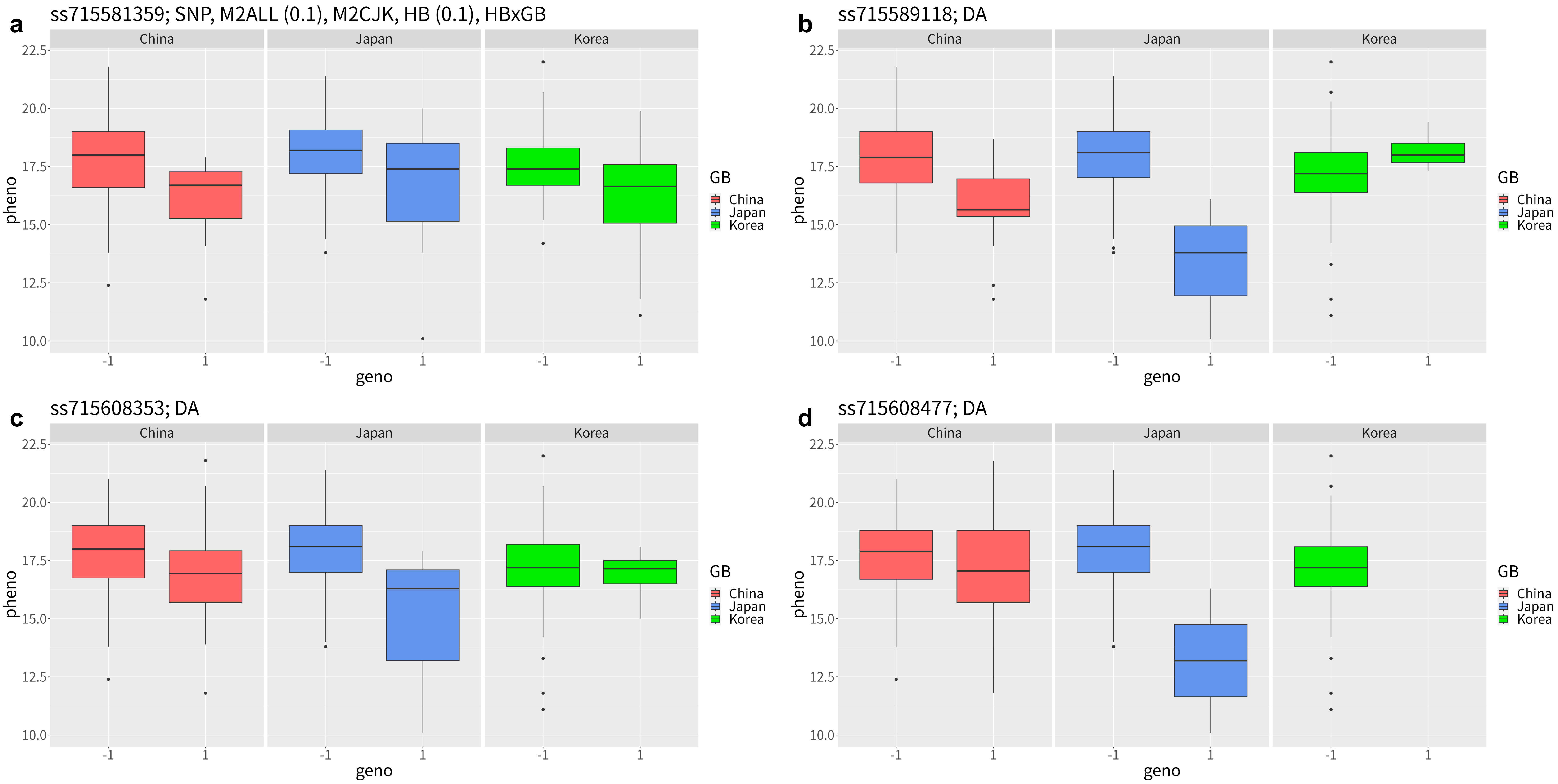


#### (2) Protein content

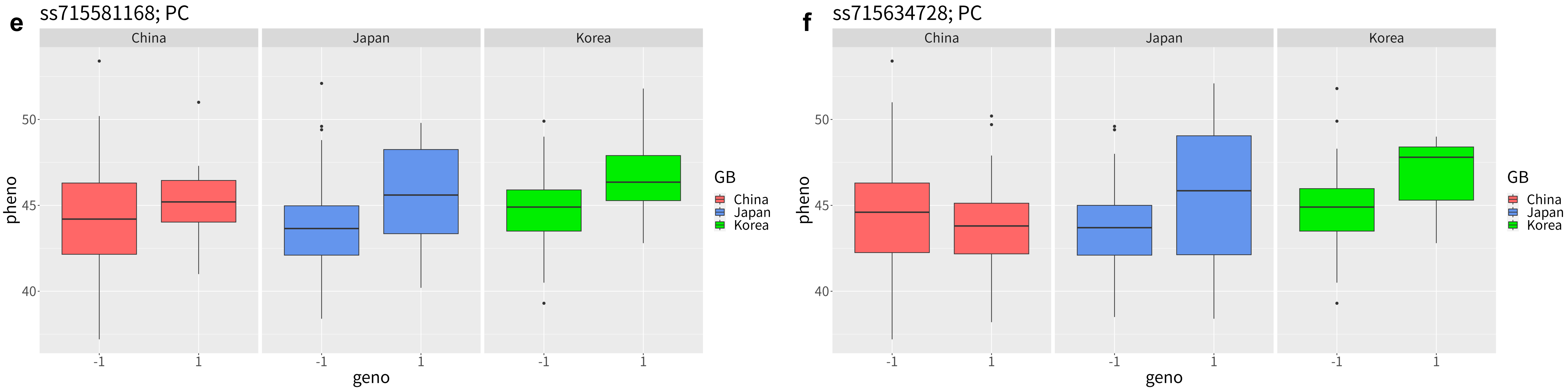
