## Supplementary material for "Development of a novel GWAS method to detect QTL effects interacting with the discrete and continuous population structure": S3 Fig

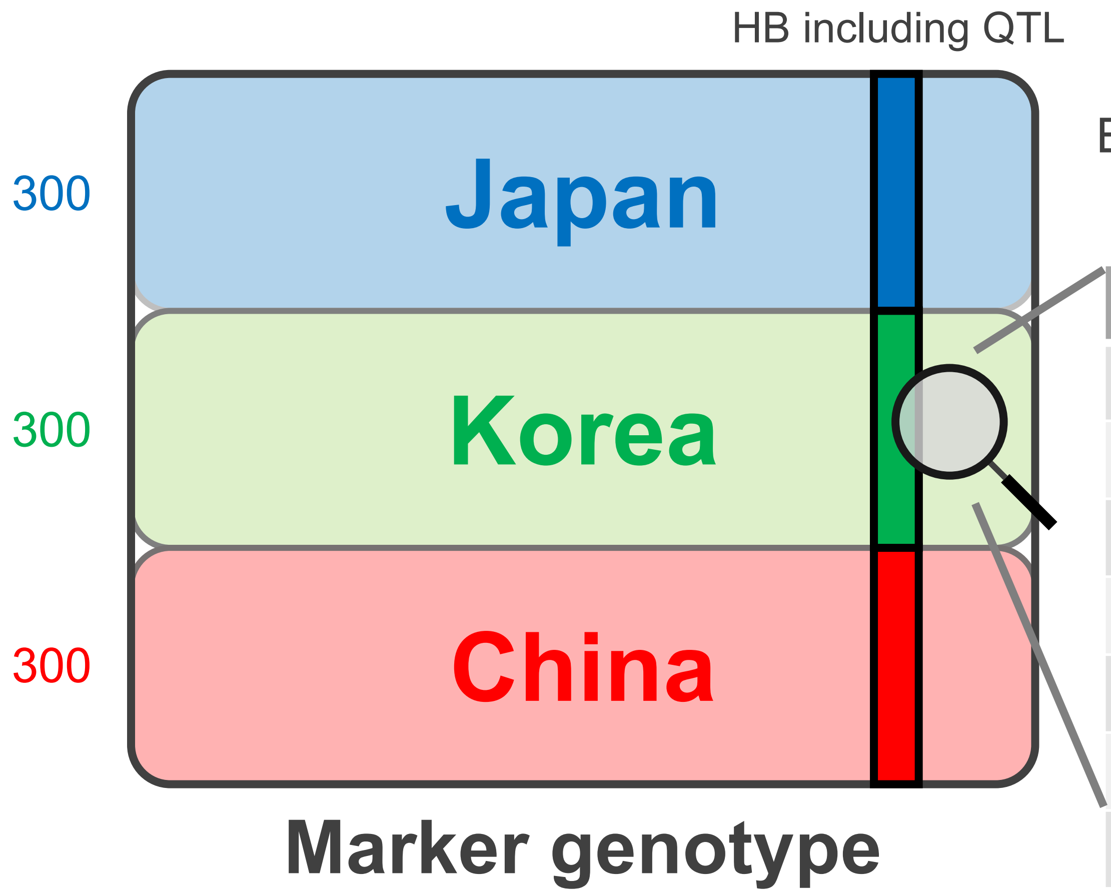


Ex.) China-specific QTL effects depending on haplotypes

(In a case where allele G in haplotype 2 & 3 has some effects.)

| | Allele | Haplotype | Country | Effect |
|---|---|---|---|---|
| Ind 1 | A | 1 | Japan | × |
| Ind 2 | G | 3 | Japan | × |
| … | … | … | … | … |
| Ind 301 | G | 4 | Korea | × |
| … | … | … | … | … |
| Ind 899 | G | 2 | China | ○ |
| Ind 900 | G | 4 | China | × |
